# Resolving power as a measure of protein stability prediction

**DOI:** 10.64898/2026.09.18.752669

**Authors:** Michele Ronco

**Affiliations:** European Commission, Joint Research Centre, Ispra, 21027, Italy

## Abstract

Rank correlation and discriminative resolution are distinct properties of a stability predictor, yet current benchmarks measure only the first. Here we introduce resolving power: the ΔΔ*G* separation at which a predictor orders two variants correctly three times in four. Applied to BioEmu across twelve wild-type domains and three supervised regressors on an independent twenty-seven, resolving powers cluster near 1 kcal mol^−1^ — comparable to the median variant-pair separation the benchmarks ask these models to rank, so most pairs cannot be ordered. A closed form ties this resolution to the calibration slope *α*; Spearman correlations account for only 31% of its variance. BioEmu extracts Δ*G* from the ratio of folded and unfolded populations, so what it samples of each becomes a question. Its folded populations reproduce experimental radii of gyration within half an ångström, whereas those it labels unfolded are collapsed chains roughly 20 to 40% below a sequence-specific random-coil reference. The ensemble still carries variant differences on panels where the sampled unfolded fraction approaches zero, but the log-odds counting readout runs out of range. Scored by a learned energy, those same conformations recover resolution on ten of twelve panels, with the gain separated from noise on four, without retraining.

## 1 Introduction

Whether an amino-acid substitution destabilises a protein, and by how much, has to be answered in order to interpret variants of unknown significance in disease-associated genes [1, 2], to advance therapeutic proteins from candidates to clinical products [6, 7], and to guide the directed evolution of new function [8–10]. In ClinVar, roughly half of pathogenic missense variants act by destabilising the fold rather than by disrupting an active site directly, whether counted by structure-based prediction across the whole proteome or by direct measurement of protein abundance in large mutagenesis experiments [3]. It is a question that resists a single-structure answer.

Variant stability is a hard target for current protein foundation models. Folding stability is not a property of the native structure but a balance between the folded ensemble and a heterogeneous unfolded one [41], so a predictor that emits a single conformation per sequence has no direct handle on the ratio the measurement reports. And the training regime that made those models accurate on folds works against variant sensitivity: pretraining on the Protein Data Bank teaches the network to reconstruct native basins, and evolutionary information collapses similar residues into shared structural output, so isolated substitutions perturb the prediction only weakly. AlphaFold2 [11], its successors [12, 13] and structure-aware protein language models recover experimental folds to within an ångström but return destabilising mutants within 0.1 to 0.6 Å of the wild type at unchanged confidence [14], acting as knowledge-based predictors of the native basin rather than models of the physics of folding [39].

Three lines of work have tried to close this gap. Sequence-only variant scorers such as ESM-1v [50] and AlphaMissense [51] respond to point mutations but predict evolutionary or clinical signal, not thermodynamic stability [3, 30]. A second line trains directly on stability: IFUM models the unfolded state as a Flory random coil [20], SaProtΔΔ*G* and Aug_ESM3ΔΔ*G* fine-tune sequence-and-structure networks on the Mega-scale corpus [22], and knowledge-based ProTherm lookups work without structural input [24]. A third generates conformational ensembles: flow-matching [16] and denoising diffusion [15, 17] models produce many conformations per sequence, fine-tuned on molecular-dynamics trajectories to approximate the distribution MD would produce.

Extracting a per-variant Δ*G* from an ensemble is a natural fit for what a substitution biophysically does. A mutation may redistribute populations across native basins whose individual structures are little altered [21, 52, 53], widen the folded ensemble without opening new states, or generate partially unfolded alternatives that reshape the free-energy landscape [54]; folding stability Δ*G* integrates over these mechanisms into a single scalar, which a single-conformation predictor cannot see and an ensemble at least admits. Among ensemble emulators we focus on BioEmu, which trains a denoising diffusion sampler on the AlphaFold Database and hundreds of milliseconds of aggregate molecular dynamics, then adds a third training stage, property-prediction fine-tuning, that reweights its ensemble against over 500,000 mutant sequences from the Mega-scale corpus [15, 18] so that experimental Δ*G* enters the objective directly. On this benchmark BioEmu is reported to reach mean absolute errors below 1 kcal mol^−1^ and Spearman correlations above 0.6 [15]. Its default per-variant Δ*G* is a counting readout: the fraction of sampled conformations classified as folded, converted to a free energy through a Boltzmann relation. As noted by the ProteinEBM authors [29], this class of ensemble emulator lacks an explicit energy function, so no principled criterion within the sampler identifies which of its generated conformations are energetically favourable, an observation that motivates the alternative readout we test below.

In this study we ask how well a stability predictor can order two variants by folding stability. Resolving power *s*^∗^ is defined as the pairwise separation at which a predictor is correct three times in four, in the units of the measurement; a closed form factorises it into a calibration slope *α* and an ensemble noise scale *σ*. Prior benchmarks have found the calibration slope smaller than 1 across many stability predictors, a compression of predictions toward zero [25]. Here we apply *s*^∗^ to BioEmu on twelve wild-type panels and to three supervised regressors on an independent twenty-seven, chosen so that their wild types sit near their unfolding transition and both stabilising and destabilising variants populate the dynamic range: seven small monomers from Mega-scale [18], three binding domains with complete deep mutational maps [5, 30], and two larger single-domain proteins with literature stability measurements [48, 49]. On the same panels, BioEmu’s folded and unfolded populations are characterised against sequence-specific random-coil references [33–35], identifying where the log-odds counting readout runs out of range though the ensemble still carries variant differences. As an alternative readout, we rank the sampled conformations by ProteinEBM [29], a sequence-conditioned energy trained on physical data, and test whether this recovers resolution where counting fails. Both the resolving power and the scoring alternative are computed from arrays that existing benchmarks already release.

## 2 Results

### 2.1 A resolving power for stability prediction

For pairs of variants of the same wild-type separated by *s* in kcal mol^−1^, the resolving power *s*^∗^ is the smallest *s* at which the predictor orders 75% of such pairs correctly [26, 27]. The quantity is invariant under monotone rescaling of the predictor’s output and expressed in the units of the measurement, so it can be compared directly with the typical variant separations of a benchmark. Table 1 summarises the twelve panels analysed and their per-panel numbers.

**Table 1:** The twelve wild-type panels. Source: cDNA display proteolysis on Mega-scale [18], abundancePCA on binding domains [30], or literature ΔΔ*G* [48, 49]. *α*: BioEmu’s calibration slope on the panel. 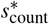 and 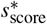: resolving power, in kcal mol^−1^, from the population-counting readout and from ProteinEBM scoring of the same conformations [29]. Sub-/superscripts: 95% percentile bootstrap intervals over variants. Dagger (†): scoring resolves better than counting at 95% paired significance (Methods). Asterisk (^∗^): wild-type present in the Mega-scale corpus, an upper bound on BioEmu’s PPFT training exposure. Dashes: not attained.

| Protein | $L$ | $n$ | Source | $\alpha$ | $s^*_{\text{count}}$ | $s^*_{\text{score}}$ |
| --- | --- | --- | --- | --- | --- | --- |
| EHEE_rd2_0152* | 40 | 100 | Mega-scale [18] | 0.48<sup>0.55</sup><sub>0.40</sub> | 0.53<sup>0.69</sup><sub>0.41</sub> | 0.54<sup>0.66</sup><sub>0.42</sub> |
| HHH_rd4_0870* | 43 | 100 | Mega-scale [18] | 0.75<sup>0.85</sup><sub>0.66</sub> | 0.76<sup>1.01</sup><sub>0.52</sub> | 0.59<sup>0.93</sup><sub>0.43</sub> |
| EA_run7_1025_0001* | 47 | 100 | Mega-scale [18] | 0.42<sup>0.62</sup><sub>0.27</sub> | 1.62<sup>2.20</sup><sub>0.68</sub> | 1.72<sup>2.23</sup><sub>0.90</sub> |
| 2CJJ* | 54 | 100 | Mega-scale [18] | 0.17<sup>0.21</sup><sub>0.12</sub> | 1.13<sup>1.57</sup><sub>0.75</sub> | 0.97<sup>1.46</sup><sub>0.64</sub> † |
| 2ZW1_V55S* | 57 | 100 | Mega-scale [18] | 0.34<sup>0.41</sup><sub>0.27</sub> | 1.03<sup>1.36</sup><sub>0.74</sub> | 0.91<sup>1.23</sup><sub>0.61</sub> |
| 2K5H* | 62 | 100 | Mega-scale [18] | 0.40<sup>0.44</sup><sub>0.35</sub> | 1.10<sup>1.57</sup><sub>0.74</sub> | 0.88<sup>1.31</sup><sub>0.59</sub> |
| 2QFF* | 70 | 100 | Mega-scale [18] | 0.16<sup>0.20</sup><sub>0.11</sub> | 0.83<sup>1.37</sup><sub>0.60</sub> | 0.80<sup>1.29</sup><sub>0.59</sub> |
| GB1* | 56 | 37 | abundancePCA [30] | 0.39<sup>0.51</sup><sub>0.26</sub> | 1.11<sup>2.06</sup><sub>0.50</sub> | 1.02<sup>1.87</sup><sub>0.45</sub> † |
| GRB2-SH3 | 56 | 21 | abundancePCA [30] | 0.46<sup>0.59</sup><sub>0.19</sub> | 0.86<sup>2.47</sup><sub>0.41</sub> | 0.59<sup>1.04</sup><sub>0.32</sub> |
| PSD95-PDZ3 | 84 | 33 | abundancePCA [30] | 0.26<sup>0.59</sup><sub>−0.01</sub> | 2.25<sup>17.55</sup><sub>0.75</sub> | 0.47<sup>0.66</sup><sub>0.28</sub> † |
| p53 DBD | 194 | 9 | literature [49] | −0.01<sup>0.06</sup><sub>−0.07</sub> | — | 1.50<sup>10.88</sup><sub>0.41</sub> |
| T4 lysozyme | 164 | 7 | literature [48] | 0.13<sup>0.17</sup><sub>−0.06</sub> | 2.02<sup>3.05</sup><sub>0.59</sub> | 0.20<sup>0.33</sup><sub>0.01</sub> † |

We illustrate the measurement on 2QFF, a 70-residue three-helix bundle drawn from BioEmu’s own folding-stability benchmark [15]. On this domain, the model produces predictions whose Spearman correlation with experiment is 0.62 and whose mean absolute error is 0.74 kcal mol^−1^ (Fig. 1a). Both numbers are typical of the model’s reported performance on this benchmark. On the same predictions the resolving power is 0.83 kcal mol^−1^, and 53% of the mutant pairs in the panel are separated by less than that (Fig. 1a). The correlation and the mean absolute error do not carry this information. Neither is expressed on the scale at which two variants would have to differ before the model can be trusted to order them, and neither is easy to compare across panels of different composition.

**Figure 1.**
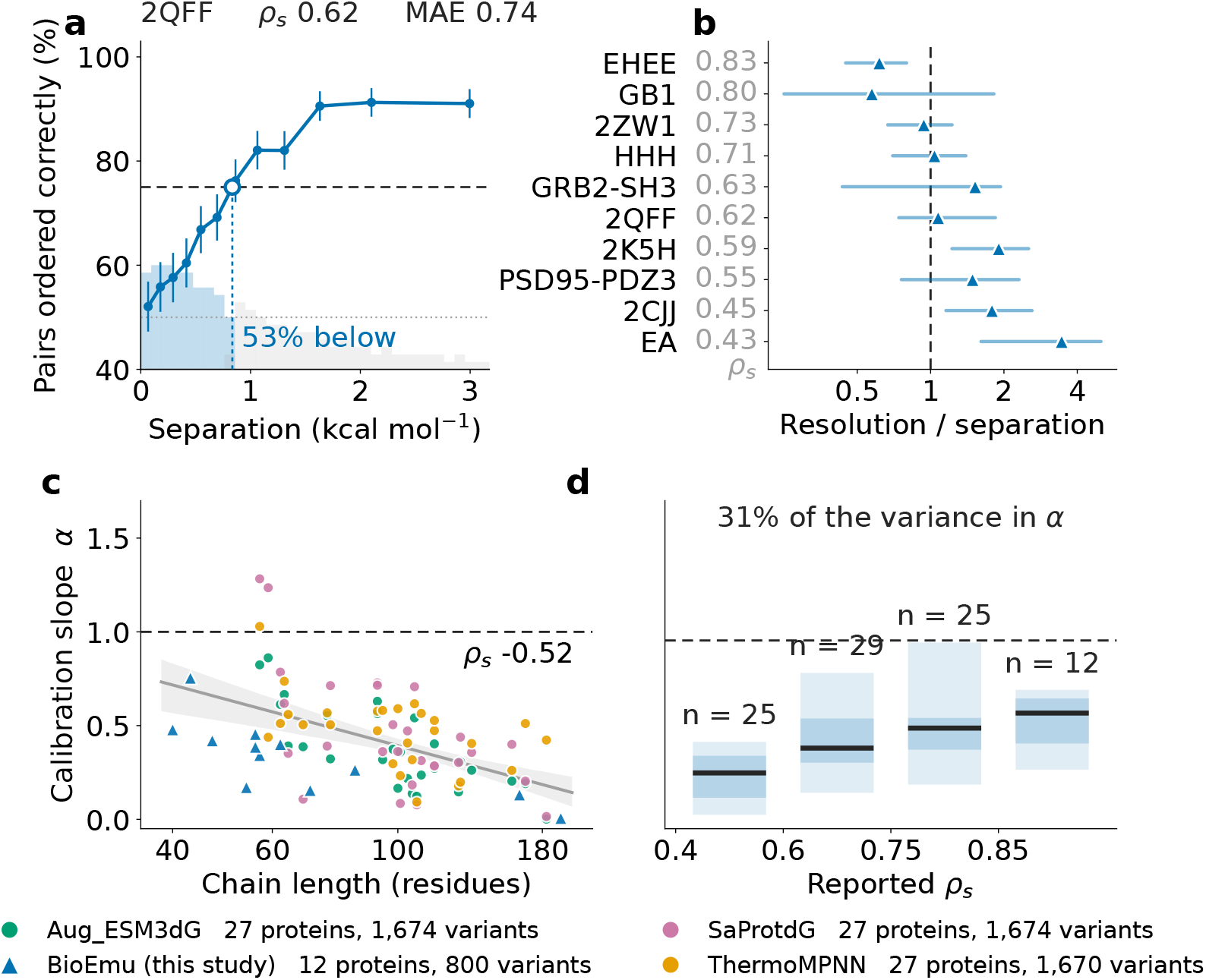
A resolving power for stability prediction. **a**, BioEmu on 2QFF: fraction of pairs ordered correctly against pairwise separation *s*. Dashed, 75% threshold; dotted, *s*^∗^; blue histogram, pairs below *s*^∗^. **b**, Resolving power divided by median pairwise separation across ten panels [18, 30], ordered by BioEmu’s Spearman correlation (grey). **c**, Calibration slope *α* against chain length for BioEmu and three regressors on S1724 [22]. Grey line and band, pooled linear fit with 95% CI. **d**, *α* distribution within four bins of reported Spearman. Line, median; bands, IQR and 5–95%. Dashed at *α* = 1 in c, d. 95% intervals throughout a and b.

Across the ten wild-types with deep mutational data (Fig. 1b), the resolving power exceeds the median pairwise separation of the same panel in seven cases, and in three of these the 95% bootstrap interval on the ratio excludes 1. Ordering wild-types by BioEmu’s reported Spearman correlation shows this is not confined to poorly ranked panels: at *ρ*_*s*_ = 0.71 the resolving power already exceeds the typical pair separation. Pooling BioEmu’s predictions across the distributed benchmark of 267 mutants from 90 wild-types gives a resolving power of 1.10 kcal mol^−1^ against a median pair separation of 0.91 kcal mol^−1^, so most pairs the benchmark asks the model to rank cannot be ordered. T4 lysozyme and p53 DBD are outside Fig. 1b at 7 and 9 mutants; 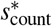 is 2.02 kcal mol^−1^ and unattained respectively (Table 1).

The resolving power has a closed form. Modelling the prediction for a mutant of true effect *x* as *f* = *αx* + *n*, with *n* Gaussian noise of variance *σ* ^2^ independent of *x*, the difference of two predicted effects for a pair separated by *s* is Gaussian with mean *αs* and variance 2*σ* ^2^. The probability of correctly ordering the pair is

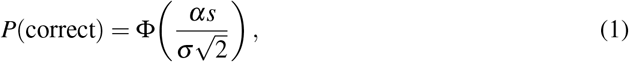

where Φ is the standard normal cumulative distribution function. Setting *P*(correct) = 3*/*4 gives the three-in-four threshold *s*^∗^,

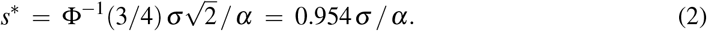

The three-in-four threshold sits midway between chance and certainty, the standard discrimination criterion in the pairwise-comparison literature [26, 27]. Related statistics average this ordering probability over the panel’s own distribution of separations rather than fixing it at one point [28]. Unlike a rank correlation, the resolving power depends only on two intrinsic properties of the predictor: the calibration slope *α* and the noise scale *σ*. Although *α* is itself attenuated by measurement error in the true effects, so a panel of narrow dynamic range returns a smaller *α* for the same predictor, *σ* is attenuated in step and the ratio *σ/α* is preserved. Across the seven proteolysis panels, *α* correlates with the panel’s measured ΔΔ*G* range at Spearman +0.89 while *s*^∗^ does so at +0.04: the invariance holds empirically on the data reported here. Spearman correlation and mean absolute error do not have this property: on a wider panel more pairs sit far enough apart to be easy to order (higher Spearman), and prediction errors scale with the effect magnitudes (larger MAE). Since *s*^∗^ scales as 1*/α*, halving *α* doubles *s*^∗^, and rank correlation is blind to this since it measures order, not scale. Consider two noiseless predictors, one with *α* = 1 and one with *α* = 0.1. Both rank the variants perfectly, so both would score Spearman = 1 in the absence of noise. Yet once the same noise *σ* is added, the second predictor’s predicted differences lie a factor of ten closer to zero relative to 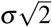, so pairs need ten times the true separation before the sign of *f*_*i*_ − *f* _*j*_ agrees with the sign of *x*_*i*_ − *x* _*j*_ three times in four. Same ranking, ten times worse resolution.

Across the twelve panels of Table 1, BioEmu’s fitted calibration slope *α* is below 1 (Fig. S1). The 95% interval on *α* includes zero on PSD95-PDZ3, T4 lysozyme and p53 DBD. To ask whether this compression is peculiar to reading a free energy off a sampled ensemble, we recomputed *α* from the released per-mutant predictions of three supervised ΔΔ*G* regressors, SaProtΔG, augmented ESM3ΔG [22, 32] and ThermoMPNN [31], evaluated on 27 wild-types of the S1724 benchmark [22] (Methods). The calibration slope is below 1 on 78 of the 81 fits (range 0.001 to 1.283), and *α* declines with chain length in all four, at pooled Spearman −0.52 (Fig. 1c). Both routes to Δ*G* here are trained on experimental stabilities, so compression is a shared property of the four current stability predictors tested [25], not specific to reading Δ*G* off a sampled ensemble.

Keeping the predictor’s *α* and *σ* fixed while increasing the standard deviation of true variant effects from 0.6 to 2.0 kcal mol^−1^ raises Spearman from 0.24 to 0.66, without any change in model performance. The four models tested show the same signature. Binning the 91 model-panel fits by reported Spearman, *α* spans a factor of 6 to 9 within a bin and adjacent bins overlap almost entirely (Fig. 1d). Reported Spearman accounts for 31% of the variance in the calibration slope *α*, and two panels at the same Spearman can differ fourfold in the fraction of a true mutational effect the model registers. Across the four predictors, resolving powers cluster near 1 kcal mol^−1^, with a supervised median of 0.99 kcal mol^−1^ across 74 fits (range 0.44 to 4.28), and on 31 of these *s*^∗^ exceeds the panel’s median pair separation, so most such pairs cannot be ordered reliably. Both the calibration slope *α* and the resolving power *s*^∗^ can be computed from arrays existing benchmarks already release, and we report both alongside every rank correlation quoted here.

BioEmu’s Δ*G* is not fit directly. It is counted from the folded and unfolded populations of a sampled ensemble, so a compression in the returned free energy could originate in what those populations contain, or in how they respond to substitution, or both. We turn to the geometry of the two states next.

### 2.2 The geometry of BioEmu’s folded and unfolded ensembles

Every conformation of every variant was classified as folded or unfolded according to its fraction of native contacts, using the same criterion BioEmu uses internally to derive Δ*G*. Radii of gyration, secondary-structure content and inter-residue contacts were then pooled per protein across all variants of that protein.

Fig. 2 covers the nine panels for which BioEmu samples enough unfolded conformations to characterise both classes (Fig. S2 for the raw joint distribution across all twelve); on the other three (PSD95-PDZ3, T4 lysozyme, p53 DBD) it does not, a finding revisited in §2.3. Across the nine, the folded ensembles reproduce their experimental structures closely. Radii of gyration agree with the crystal reference to within 0.5 å on all nine, at a median deviation of 0.11 å across chain lengths from 40 to 70 residues (Fig. 2a, green). Secondary-structure content is lower than the crystal value by a uniform 8 to 13 percentage points. Because the same frames, classification and radius definition recover a quantity determined independently by X-ray crystallography, this provides a control for the measurements that follow.

**Figure 2.**
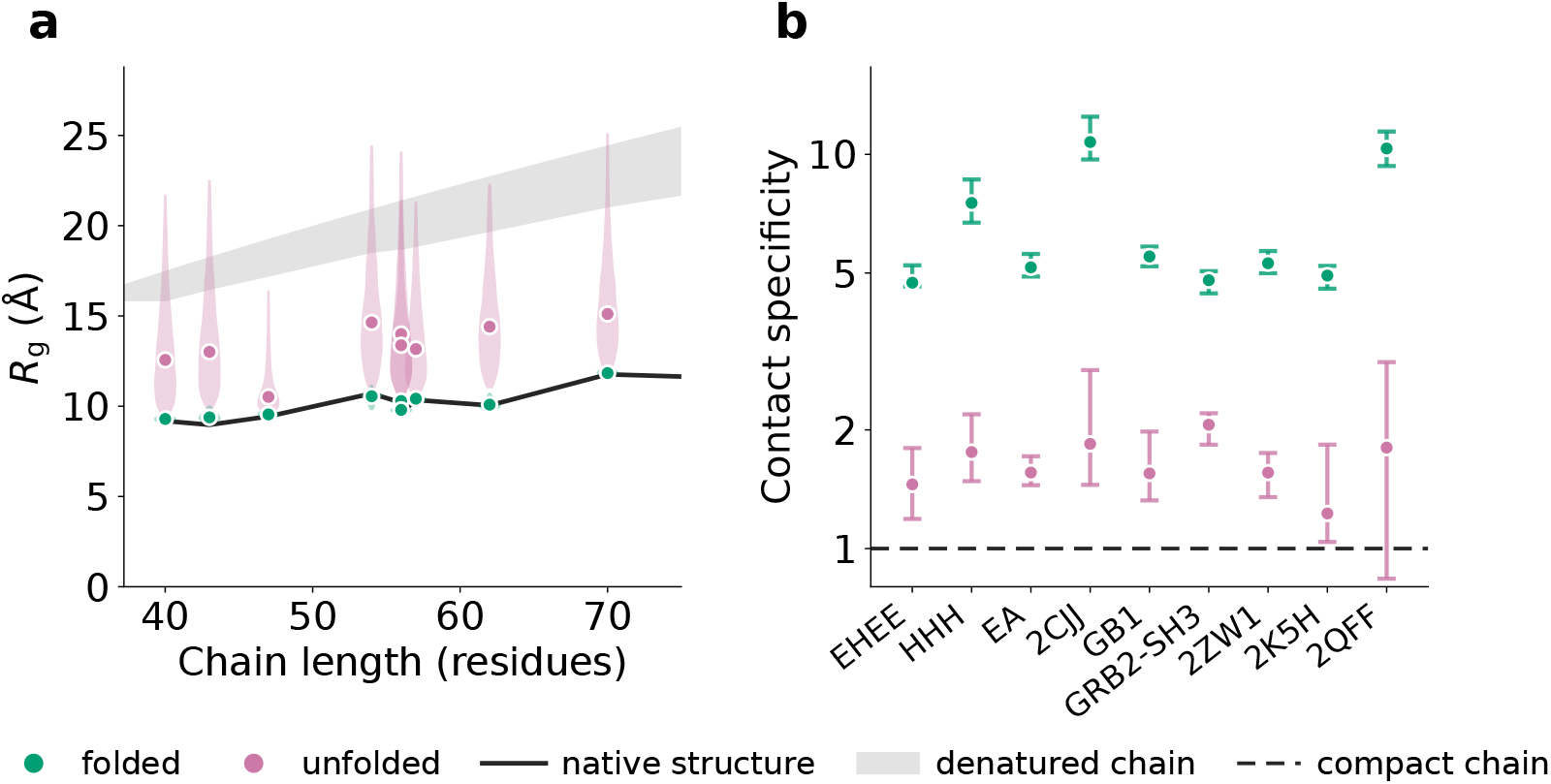
Geometry of folded and unfolded ensembles across nine benchmark domains of chain length up to 75 residues. **a**, Radius of gyration *R*_*g*_ against chain length. Green points, median *R*_*g*_ of the folded ensemble; pink violins, distribution of *R*_*g*_ in the unfolded ensemble; solid black line, *R*_*g*_ of the crystal structure; grey band, reference for a denatured chain of the same length, bounded above by an empirical Flory scaling [36] fitted to scattering on 28 chemically denatured proteins and below by the sequence-specific Analytical Flory Random Coil expectation [33]. **b**, Contact specificity, the ratio of the fraction of long-range native contacts (sequence separation *>* 8) in each ensemble to the same fraction in a self-avoiding chain grown to the ensemble’s measured *R*_*g*_. Green, folded; pink, unfolded. Dashed line at 1, the compaction-matched baseline. Points, median across variants pooled per domain; bars, 25th to 75th percentile.

The unfolded ensembles do not, and the manner of the failure is not a simple shift. Their radius-of-gyration distributions are broad: standard deviations reach 1.9 to 3.6 å, an order of magnitude wider than the 0.10 to 0.33 å of the folded ensembles, and their upper tails reach the dimensions a denatured chain of the same length should have. Their centres do not. Median radii lie at 0.55 to 0.73 of an empirical Flory scaling fitted to single-molecule scattering on 28 chemically denatured proteins [36], and at 0.62 to 0.81 of the sequence-specific Analytical Flory Random Coil expectation [33] (Fig. 2a, purple; the grey reference band is bounded above by the former and below by the latter; per-panel values in Table S2). The model can generate expanded chains. It generates too few of them. This breadth is a property of each variant’s own ensemble rather than of the mutations, since decomposing the pooled variance leaves 93 to 99% of it within variants.

Compaction alone does not distinguish a slackened native structure from a collapsed chain. The first would retain its native contacts, the second would make contacts by proximity. We therefore counted, for each ensemble, the fraction of long-range native contacts (sequence separation greater than eight residues) it makes, and compared it to the same fraction in a self-avoiding chain grown, not rescaled, to the same measured radius of gyration. Rescaling would shorten the bonds and inflate the comparison. Folded ensembles exceed this compaction-matched baseline by 4.7- to 10.7-fold. Unfolded ensembles exceed it by only 1.2- to 2.1-fold (Fig. 2b). The restriction to long-range pairs is necessary. Including local pairs makes the excess correlate with residual secondary structure at *ρ* = +0.86, an association that reflects a helix placing residue *i* within contact range of *i* + 4 rather than any tertiary organisation. Once such pairs are excluded the correlation falls to *ρ* = +0.11. The conformations the model labels unfolded are therefore collapsed chains of the wrong dimensions rather than loosened folds. That distinction separates a solvation-like error from a missing conformational class, and it matters because the two admit different remedies.

A reference state should also be insensitive to single substitutions. Across the roughly one hundred variants of each domain, the sequence-specific coil expectation varies by 0.02 to 0.04 å, whereas the median unfolded radius shifts by 0.66 to 2.25 å, between 20- and 68-fold more. Allowing for sampling error in the per-variant means, the discrepancy remains at least an order of magnitude. Taken together with the dimensions and the contact composition, the picture is consistent. The ensembles labelled unfolded are not random coils, so a coil expectation neither describes their size nor bounds their response to mutation.

We do not claim that the geometry documented here explains the compression of Fig. 1 — the compaction is uniform across proteins whose calibration slopes vary fivefold, and the three supervised regressors of Fig. 1c compress by the same amount without modelling an unfolded state at all — but Fig. 2 does establish that what BioEmu treats as an unfolded reference is not the reference state a folding free energy requires. We therefore ask whether the same conformations can be scored differently, without touching the pipeline.

### 2.3 Scoring the sampled ensemble recovers resolution where counting fails

Assigning an energy to each conformation avoids the need for an unfolded reference. We scored every conformation in each BioEmu ensemble with ProteinEBM [29], an energy-based model trained by score matching on structural databases and molecular-dynamics trajectories, and took the ensemble mean as the per-variant readout (Fig. S3 for the per-panel folded/unfolded energy distributions). No threshold, no folded-unfolded population ratio, no separately constructed reference. This also bypasses the unfolded-state approximation used by ProteinEBM’s own ΔΔ*G* estimator, which constructs a Ramachandran-random reference. On PSD95-PDZ3 (Fig. 3a), two variants whose measured effects differ by 1.11 kcal mol^−1^ are separated by 0.04 standard deviations under BioEmu’s counting readout and by 8.1 standard deviations under scoring, measured on each readout’s spread across sixty independent ensembles per variant. Two peaks are resolvable in the Rayleigh sense once their separation exceeds their combined spread; scoring resolves this pair, counting does not. The panel’s resolving power improves from 2.25 to 0.47 kcal mol^−1^.

**Figure 3.**
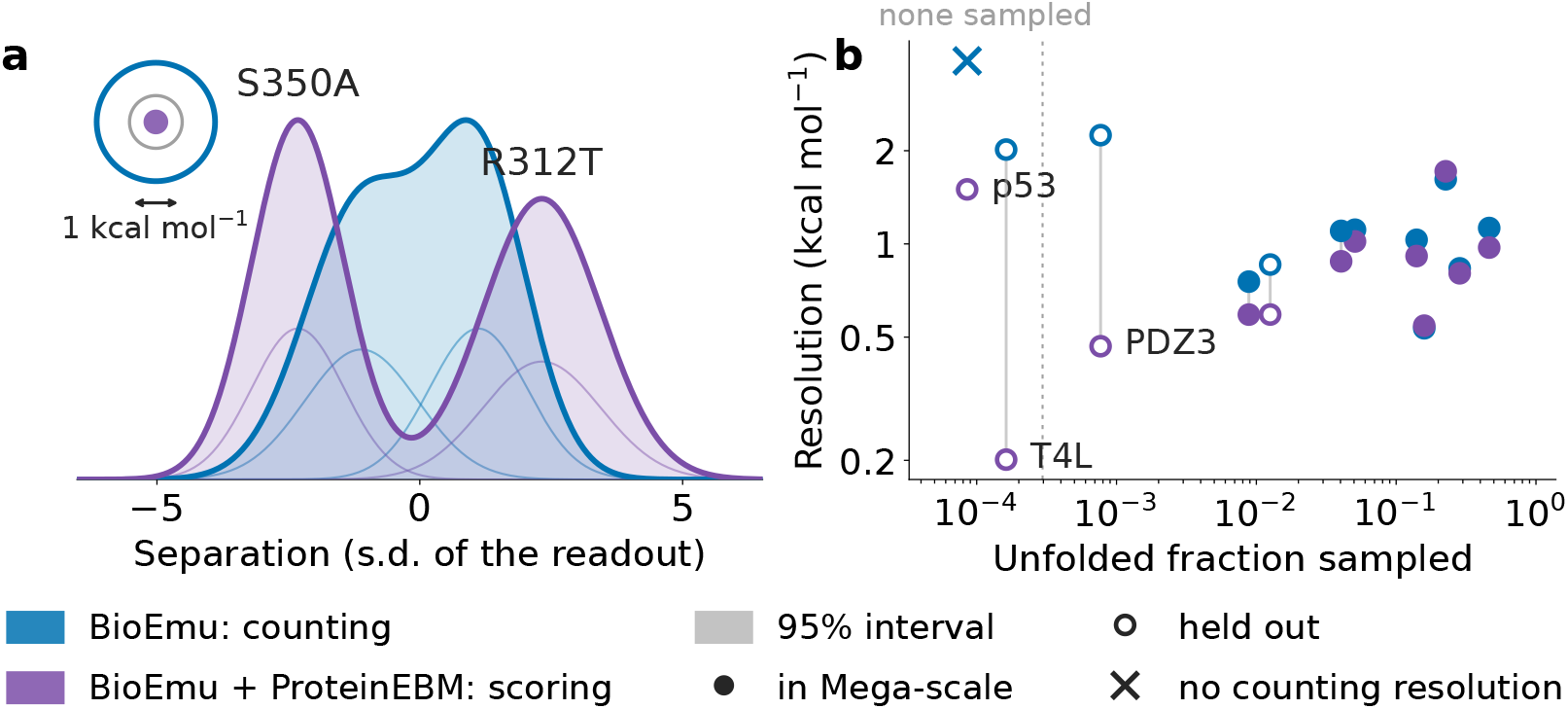
Scoring the sampled ensemble recovers the panels where counting fails. **a**, Distributions of BioEmu’s two readouts on PSD95-PDZ3 across sixty independent ensembles per variant, for two mutants (S350A, R312T) whose measured effects differ by 1.11 kcal mol^−1^. Under counting the variant distributions overlap (0.04 standard deviations apart, blue); under scoring they separate (8.1 standard deviations apart, purple). Lighter shading, per-variant density; heavier outline, sum. Inset, a 1 kcal mol^−1^ interval at scale on each readout’s axis. **b**, Resolving power against BioEmu’s sampled unfolded fraction across the twelve panels. Circles, counting (blue) and scoring (purple), connected by a line per panel. Filled, panels whose wild-type is in the Mega-scale dataset (Methods); open, not present. Cross, no counting resolution attained (p53 DBD).

Across the twelve panels the gain is conditional (Fig. 3b). Scoring resolves better on ten of twelve panels by point estimate, with the improvement exceeding the paired 95% bootstrap interval on four: 2CJJ, GB1, PSD95-PDZ3 and T4 lysozyme (Table 1, daggers). The largest gains are on the panels where BioEmu samples the smallest unfolded fraction, the regime in which the counting readout saturates. The folded-fraction estimator hits its ceiling and the log-odds Δ*G* loses its signal. On p53 DBD and T4 lysozyme the model samples essentially none, so no counting resolving power exists to improve on, and scoring returns 1.50 and 0.20 kcal mol^−1^ respectively. Nor are the populations Boltzmann-weighted where BioEmu does sample the unfolded state: on the seven proteolysis panels the sampled unfolded fraction exceeds the equilibrium expectation from the measured Δ*G* by 4 to 119-fold, while the three abundance panels are under-populated. But improvement is not confined to that regime. 2CJJ provides a counterexample: BioEmu samples 46% unfolded conformations on that panel, the highest of the twelve, yet scoring still resolves better than counting at paired significance. Because the conformations are unchanged, the gain cannot arise from improved sampling; it arises from how the sampled ensemble is read out. The ensemble does respond to substitutions and weakly tracks effect magnitude on most panels (Fig. S4, Table S1). Subsampling the seven proteolysis panels to the 7 to 33 mutants of the smaller panels leaves the gap between readouts unchanged in sign, ruling out a small-sample artefact. Scoring improves discrimination on this benchmark, not calibration. The energies are returned in arbitrary units, the calibration slope *α* is unchanged, and the compression documented in §2.1 is not addressed by scoring alone.

An alanine scan localises the signal on the substituted residue. Replacing each position in turn by alanine, in both the variant and the wild-type sequence, shifts the prediction most at the substituted site on 86 to 94 per cent of variants; scaling each variant’s profile by its own largest shift, the median elsewhere is 0.18. Selecting conformations by fraction of native contacts rather than at random, at matched subset sizes, does not improve resolution on any panel, with every bootstrap interval 0.5 to 0.9 kcal mol^−1^ wide, so the discrimination is spread across the ensemble rather than concentrated in its folded or unfolded conformations.

## 3 Discussion

Reading current protein stability predictors as measuring instruments, we find they resolve variants at roughly the scale on which those variants differ (median 0.99 kcal mol^−1^ across 74 supervised regressor fits, 1.10 on BioEmu pooled), so most benchmark pairs cannot be ordered. This is not specific to BioEmu, and not specific to the generative route to Δ*G*. Three supervised regressors on an independent benchmark show the same compression, a calibration slope below 1 on 78 of 81 fits that a rank correlation cannot see, and Spearman captures only 31% of its variance across the 91 model-panel fits considered here. For BioEmu, the readout compounds the effect on panels where the sampled unfolded fraction approaches zero. Its counting readout takes the log-odds of the folded fraction, which loses range as that fraction approaches 1, regardless of what the ensemble contains. Scoring the same conformations with a sequence-conditioned energy [29] recovers discrimination on ten of twelve panels, with the gain separated from noise on four (§2.3). The ensemble carries variant-dependent information that counting cannot read.

The pattern is not new to the field. The low dynamic range of ΔΔ*G* measurements in the same benchmark has been proposed as a source [22]. A natural upper bound on prediction accuracy has been placed on the basis of experimental noise alone [37]. A comparable ceiling limits protein–ligand binding free energy calculations [38]. And the general observation that current structure predictors do not learn the physics of folding, in the sense that they are largely knowledge-based predictors of the native state, has been made forcefully [39]. An independent evaluation of BioEmu on kinase mutations outside its training set reports a related observation, that mutations appear to add random noise to the ensemble rather than shift it towards the true mutant response [40]. The low-dynamic-range attribution locates the difficulty in the measurements; our low-calibration-slope diagnosis locates it complementarily in the predictor.

Where the unfolded reference state is concerned, our diagnostic converges with an emerging architectural instruction. BioEmu’s “unfolded” ensembles sit at 55–73% of the Flory scaling fitted on chemically denatured proteins [36] and at 62–81% of the sequence-specific Analytical Flory Random Coil expectation [33], with long-range native-contact composition 1.2- to 2.1-fold above a compaction-matched self-avoiding baseline. That such an unfolded reference matters for Δ*G* prediction has been shown on entirely different architectural grounds [20]: making the Flory random coil an explicit component of the network raises Mega-scale Pearson from 0.70 to 0.78 in ablation. That approach is prescriptive where ours is diagnostic. We measure how far BioEmu’s implicit unfolded state deviates from the reference IFUM was built to enforce, in a model whose training pipeline does not enforce it. The two findings converge on the same instruction: whatever the route to a folding free energy, the unfolded state has to be described rather than implied [41], a principle that physics-based Boltzmann generators were designed to respect from the outset [42]. The geometric mismatch does not by itself explain the class-level compression, since three supervised regressors that model no unfolded state at all compress by the same margin (§2.1). Within BioEmu, however, the geometric shortfall and the log-odds saturation share a common source on the saturating panels: undersampling of the unfolded state. On those panels the two failures collapse into one and cannot be cleanly separated.

Two upstream stages of BioEmu’s training pipeline are candidate causes of the compression, neither of which we test directly. AFDB pretraining teaches the model to sample near AlphaFold’s structural prediction for each input sequence, and that prediction responds weakly to point mutations [14], so sequence substitutions propagate weakly into the sampled distribution. Property-prediction fine-tuning supervises the folded-unfolded population ratio against a scalar Δ*G* without providing structural signal about which conformations should carry that free energy. The objective teaches magnitude, not redistribution. The same underpopulation silences the counting readout wherever the unfolded fraction approaches zero, so the folded-fraction estimator saturates against its ceiling and the log-odds Δ*G* loses its signal, a readout failure distinct from the geometric one that a scored readout can repair without touching the sampler.

Efforts to move protein foundation models past their PDB-centred training bias converge from different angles. Thermodynamically informed machine-learned coarse-grained potentials explicitly decompose the free energy into energetic and entropic components, guaranteeing physically consistent extrapolation across temperature regimes [44]. Latent-space optimization of AlphaFold3 embeddings maximizes ensemble log-likelihood under structural and force-field priors, sampling from their product distribution [45]. Gradient-based guidance of AlphaFold3’s reverse-diffusion trajectory with experimental likelihoods from NMR, X-ray crystallography and cryoEM data recasts the model as a sequence-conditioned prior for posterior inference over ensembles [46]. Energy-based alignment fine-tunes the sampler itself with feedback from a physical energy model [47]. What these directions share, and what our scoring layer instantiates in the simplest form, is the recognition that a PDB-trained sampler and a thermodynamically calibrated readout are two separable problems. Our approach is the least invasive of them, requiring no retraining, no experimental data at inference, and no modification of sampling. The ensemble BioEmu returns already contains the discriminative information, and extracting it more efficiently through a statistical potential is a lower bar than repairing the ensemble itself. This use of a learned energy to rerank samples from generators that lack one is proposed as a route to structure prediction and ΔΔ*G* estimation [29]; we apply it here to BioEmu ensembles and measure its effect on stability resolving power.

The calibration slope *α* and resolving power *s*^∗^ should be reported per panel alongside every rank correlation, on the same terms as the mean absolute error already is. Both are computable from arrays existing benchmarks already release, and their adoption provides a per-model, panel-independent measure of discriminative power in the units of the experimental measurement.

A targeted comparison across unfolded-state treatments would isolate whether the resolving power is bounded more tightly by reference-state error or by training-panel dynamic range. The four axes to compare are BioEmu’s implicit sampled ensemble, IFUM’s explicit Flory reference, our scoring layer, and independent BioEmu evaluations run under other downstream constraints [19, 43]. The scoring result of §2.3 already indicates that on ensemble emulators, the readout carries as much of the loss as the ensemble itself. Whether the same compression affects insertion/deletion mutations or double mutants, and whether it varies systematically across protein classes beyond those represented in the Mega-scale dataset, remain open questions.

In conclusion, the physical scale at which a stability predictor can discriminate is a first-class quantity, computable, portable across benchmarks, and expressed in the units of the measurement. Its adoption alongside the rank correlations that currently drive benchmarking would provide a more complete picture of what current tools can and cannot do, and where the next generation of predictors should aim. For clinical applications, that scale determines whether a predictor can be trusted to order the sub-kcal mol^−1^ differences separating, for instance, pathogenic from benign TP53 variants [4].

## 4 Methods

### Panels

Twelve wild-type domains were analysed, each with the set of single-substitution mutants for which an experimental ΔΔ*G* is available (Table 1). Seven come from the Mega-scale cDNA display proteolysis dataset [18]: four natural domains (2CJJ, 2K5H, 2QFF, 2ZW1_V55S_) and three *de novo* designs (EHEE_rd2_0152, HHH_rd4_0870, EA_run7_1025_0001), with about 100 mutants each. Three come from abundancePCA, the folding-stability arm of the double deep protein-fragment complementation assay [30]: GB1, GRB2-SH3 and PSD95-PDZ3, with 37, 21 and 33 mutants. Two carry ΔΔ*G* values from the primary literature: T4 lysozyme [48] and p53 DBD [49], with 7 and 9 mutants.

The seven proteolysis panels were assembled from the Mega-scale table rather than taken from BioEmu’s distributed benchmark, of which only 2QFF and 2ZW1_V55S_ are members; for those two, our ΔΔ*G* values agree with the benchmark’s to within rounding. Wild-type membership of the Mega-scale dataset was established by exact sequence match against its 477 distinct wild-type sequences, not by name, since the naming conventions of both sources admit near-collisions between different point mutants of the same parent. Mega-scale is also the source of BioEmu’s property-prediction fine-tuning (PPFT) data. The train–test split is not public, so membership of the dataset is what Table 1 records and what the asterisks denote; it is an upper bound on training exposure, not a statement of it.

### Conformational sampling

Ensembles were generated with BioEmu v1.2 (bioemu 1.4.1) using the DPM denoiser and the package’s physicality filter, from multiple sequence alignments retrieved once per wild-type and reused unchanged for every mutant of that wild-type, so that differences between variants of a panel arise from the substituted residue and not from alignment depth. Sampling depth was set per variant from the expected unfolded population, so that a stable protein receives enough draws for its unfolded state to be estimated at all: 2,000 conformations as a floor and up to 10^4^ where the expected unfolded fraction is below 10^−3^, split across ten independent seeds for wild-types and two for mutants. In total 1,470 sampling runs were performed on NVIDIA RTX PRO 4500 accelerators, yielding 3.17 × 10^6^ retained conformations.

### Readouts

#### Fraction of native contacts and the counting readout

Reference contacts were taken from the experimental structure as C*α*–C*α* pairs separated by more than three residues in sequence and less than 10 å in space. For a conformation with distances *d*_*i j*_ and reference distances 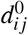, the fraction of native contacts is the soft indicator

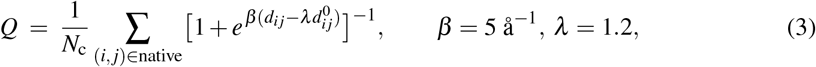

following BioEmu’s own benchmark. The folded fraction of an ensemble is the mean over conformations of a second sigmoid in *Q* centred at *Q* = 0.5, and the free energy of folding is its log-odds, Δ*G* = −*k*_B_*T* ln *p*_f_*/*(1 − *p*_f_), evaluated at *T* = 295 K, the temperature at which `bioernu-benchrnarks` performs this conversion. This is the *counting* readout throughout; ΔΔ*G* is the difference between a mutant’s Δ*G* and the wild-type’s. The equilibrium expectation for the unfolded fraction of a variant, used to compare against BioEmu’s sampled populations in §2.3, is 1*/*(1 + *e*^Δ*G/RT*^) at *T* = 295 K, and the ratio of the sampled fraction to this expectation defines the over-population factor.

#### Scoring readout

Every conformation was scored with ProteinEBM [29] (checkpoint `rnodel_6_expert_frozen_1rn_rnd`, the MD-finetuned expert model designated for scoring), at diffusion times *t* ∈ {0.01, 0.05, 0.1}; *t* = 0.1 is reported, being the value selected on validation data in [29]. The per-variant readout is the arithmetic mean of the conformation energies. The mean is the first cumulant of the quantity a free energy uses, since ln 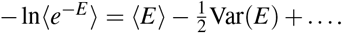 Before any panel was scored, the energy of one structure under sixteen random orientations was checked to vary by less than 2% of its mean, since the model is not equivariant by construction.

### Resolving power

For each ordered pair of mutants (*i, j*) of a wild-type, the separation is *s* = |ΔΔ*G*_*i*_ − ΔΔ*G*_*j*_| and the pair is ordered correctly when the predicted difference has the sign of the measured one; the predictor’s global orientation is taken from its own rank correlation on that panel, and ties count as incorrect. Pairs are binned into up to eight quantiles of *s*, and *s*^∗^ is obtained by linear interpolation between the two bin centres that bracket an accuracy of 3*/*4, requiring that accuracy remain above the threshold for all larger separations. The number of bins adapts to the number of pairs, keeping at least fifteen pairs per bin and never fewer than three bins, so that small panels are estimated coarsely rather than not at all; the same rule is applied to both readouts on every panel and on every subsample. The resolution can also be obtained by fitting Φ(*κs*) with a single free parameter *κ*, which is more precise on panels whose separations cluster; the binned estimate is reported throughout for both readouts, so comparisons between them are unaffected.

Under the measurement model *f* = *αx* + *n* with *n* ~ *N*(0, *σ* ^2^) independent of the true effect *x*, a pair separated by *s* has predicted difference *N*(*αs*, 2*σ* ^2^), so the probability of ordering it correctly is 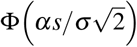, from which the three-in-four criterion gives Eq. 2. The calibration slope *α* is the least-squares slope of predicted on measured ΔΔ*G* over mutants only, the wild-type being the reference and excluded. Intervals on *s*^∗^, *α* and their differences are percentile bootstraps over variants, 2,000–4,000 resamples. Comparisons between readouts are *paired*: variants are resampled once and both readouts recomputed on the same resample, since the two share their sampling noise and independent intervals would be far too conservative. Per-bin ordering-accuracy intervals in Fig. 1a are Wilson binomial 95% intervals.

Daggers in Table 1 indicate 95% paired significance for scoring versus counting. On the seven proteolysis panels the test is the paired bootstrap of the readout difference over variants defined above. On the five abundance and literature panels, whose smaller and more heterogeneous variant samples do not support that test, the same significance is drawn from the independent-ensemble spreads of each readout on the representative pairs used in Fig. 3a.

Calibration slopes for the supervised predictors in Fig. 1c were computed from the per-mutant predictions released with ref. [22], hosted at Zenodo [23], using this same estimator, for the 27 of 35 S1724 wild-types with at least 15 mutants; below that a within-protein slope has an interval wider than the effect being measured.

### Ensemble geometry

Radii of gyration are computed over C*α* atoms and compared with the same quantity from the experimental structure. Secondary-structure content is the DSSP helix-plus-strand fraction. The denatured-chain reference band is bounded above by the Flory scaling *R*_*g*_ = 1.927 *N*^0.598^ å fitted to single-molecule scattering on 28 chemically denatured proteins [36] and below by the sequence-specific Analytical Flory Random Coil expectation [33–35], computed per variant.

To distinguish a collapsed chain from a loosened fold (Fig. 2b) we ask how many of a state’s long-range contacts the native structure also makes, relative to what compaction alone would produce. For each ensemble, we generated self-avoiding chains of the same length whose radius of gyration matches that ensemble’s measured median, by *growing* them to that radius rather than rescaling a larger chain, since rescaling shortens the virtual bonds and inflates the contact count. The specificity is the ratio of the fraction of contacts that are native in the ensemble to the same fraction in the matched random chains. Only pairs separated by more than eight residues in sequence are counted: with local pairs included, the excess tracks residual helix content at Spearman +0.86, because a helix places residue *i* within contact range of *i* + 4 irrespective of any tertiary organisation, and the association falls to +0.11 once such pairs are excluded.

### Ensemble and embedding response to substitution

For each variant we computed the Jensen–Shannon divergence (base 2, 40 bins spanning [0, 1]) between its distribution of fraction of native contacts and the wild type’s, and the first Wasserstein distance in ångströms between their radius-of-gyration distributions. A distribution redrawn from one sequence is not identical to itself, so both are reported against a floor: the same distances between two independent ensembles of the wild type, its sampling seeds split into halves of equal count, each half holding more than 200 conformations. An uneven split makes the smaller half noisier and inflates the divergence, so panels with an odd number of seeds drop one. Floors for the abundancePCA and literature panels were computed where those wild-type ensembles reside and imported; a floor from frames pooled across a panel’s variants carries between-variant variation as well as sampling variation and is not equivalent. Changes in AlphaFold2’s single and pair representations are relative Frobenius norms, ∥*X*_mut_ *X*_wt_∥ − ∥*/*∥ *X*_wt_∥, computed from the representations cached during sampling, which are keyed on sequence and alignment and are therefore the ones the ensembles were generated from. Spearman correlations against the magnitude of the measured effect are reported for panels of at least nine variants, with 95% percentile bootstrap intervals over variants.

### Repeated-ensemble spreads and ablations

The widths drawn in Fig. 3a are measured, not assumed. Blocks of conformations of the size used throughout were drawn *without replacement* within a seed and the readout recomputed, sixty times per variant; a bootstrap of a single ensemble would understate the spread of a genuinely new one. Panel a is drawn in units of each readout’s own measured spread because the two readouts have different units.

Four ablations pair each variant’s sequence with conformations other than its own: the wild-type’s ensemble, an ensemble pooled across all variants of the panel, a single experimental structure, and the variant’s own conformations under a different sequence. The last uses a *random* sequence of the same length rather than another variant’s, because two single mutants of one wild-type perturb the energy by a few units where a random sequence perturbs it by a few hundred, and a sibling control cannot fail.

## Data and code

The code for the metric and the analyses presented in the figures is available at https://github.com/mikronco/rayleigh-ddg.

## Supplementary figures

**Figure S1.**
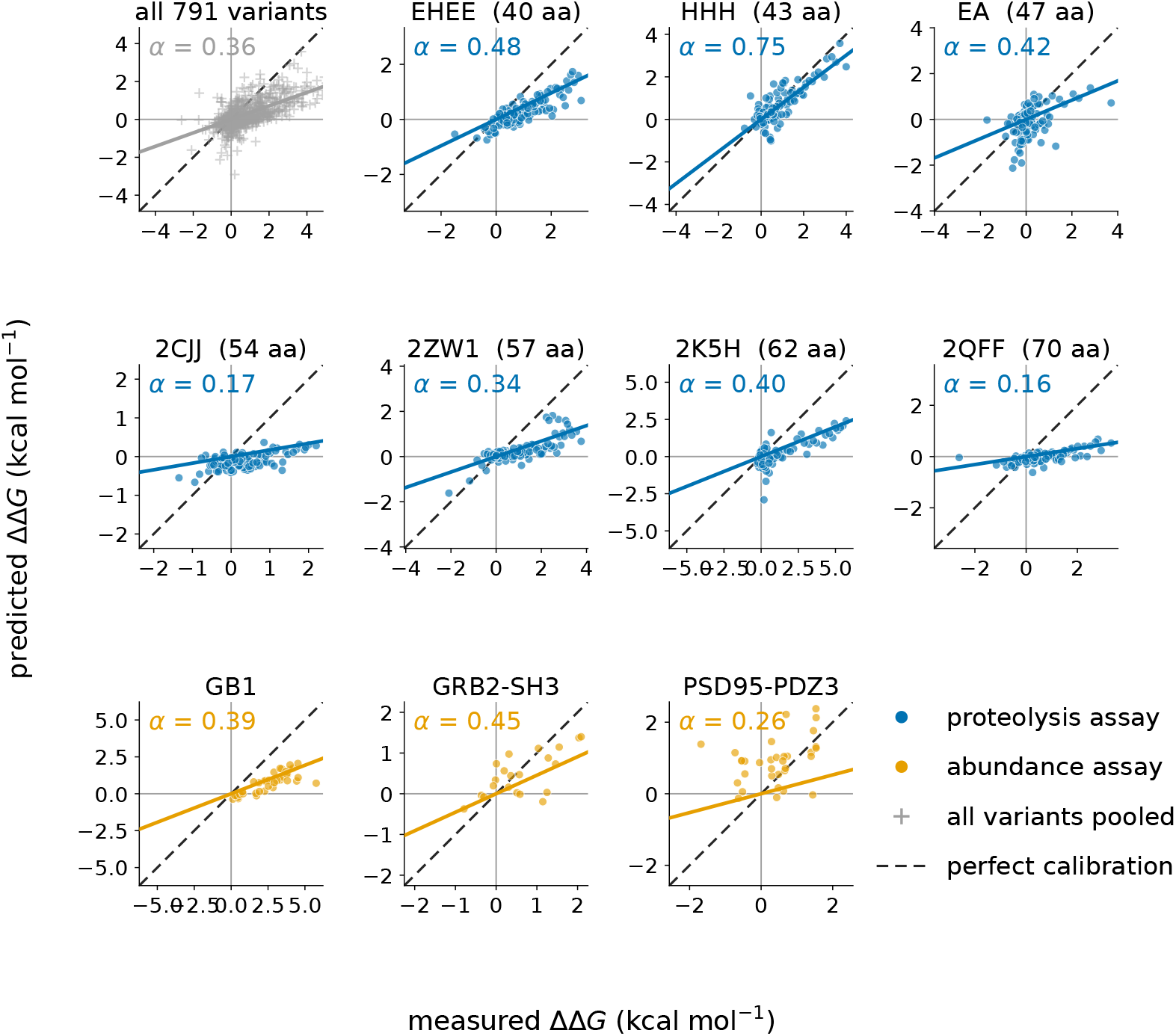
Per-panel calibration of BioEmu on the ten wild-types of Fig. 1b. Predicted against measured ΔΔ*G* for each wild-type separately, and pooled across all 791 variants (top-left panel, grey crosses; pooled *α* = 0.36). Blue circles, mutants from cDNA display proteolysis [18]; orange circles, mutants from the abundance-based fitness assay [30]. Solid coloured line, ordinary-least-squares fit of predicted on measured ΔΔ*G*; the fitted calibration slope *α* is printed above each panel. Dashed line, perfect calibration (*α* = 1). The per-panel *α* values feed the resolving-power ratios of Fig. 1b through Eq. 2.

**Figure S2.**
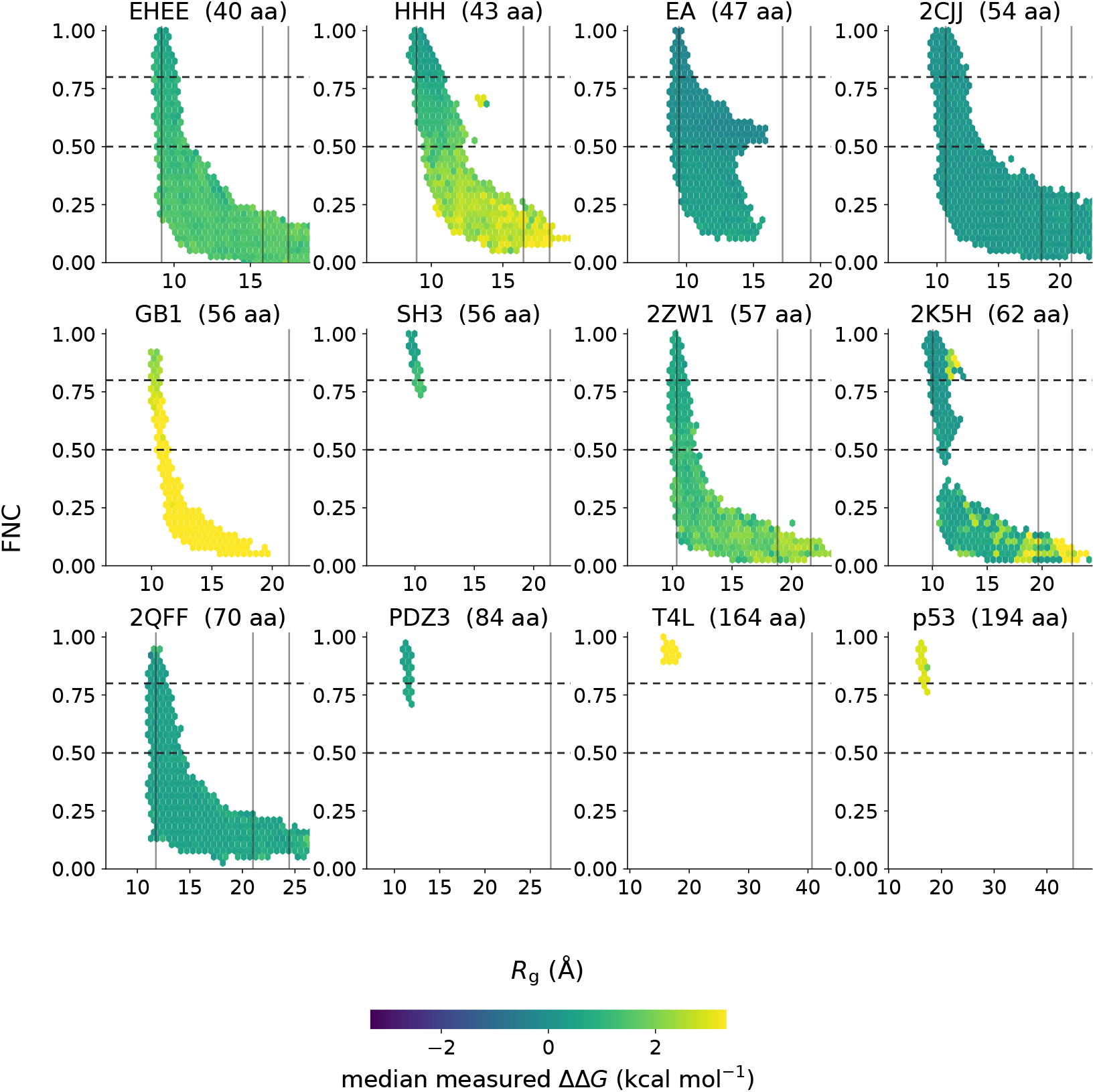
Joint distribution of fraction of native contacts and radius of gyration for all twelve wild-type panels, coloured by variant. ΔΔ*G*. For each protein, hexbins show the joint density over the (*R*_*g*_, FNC) plane of all sampled conformations pooled across all variants. Bin colour, median measured ΔΔ*G* of the variants whose conformations populate the bin (negative values stabilising, positive values destabilising). Horizontal dashed lines at FNC = 0.5 and 0.8 separate unfolded, ambiguous and folded conformations under BioEmu’s own classifier. Vertical grey lines, from left to right, the crystal-structure *R*_*g*_, the sequence-specific Analytical Flory Random Coil expectation [33], and the Hofmann-Flory expectation for a denatured chain of the same length [36]. The figure covers all twelve panels of Table 1: on the nine of Fig. 2 both classes are visibly populated, the unfolded class lying below both denatured-chain references; on the remaining three (PSD95-PDZ3, T4 lysozyme, p53 DBD) BioEmu samples almost no unfolded conformations. Destabilising mutations tilt density toward the lower-right region.

**Figure S3.**
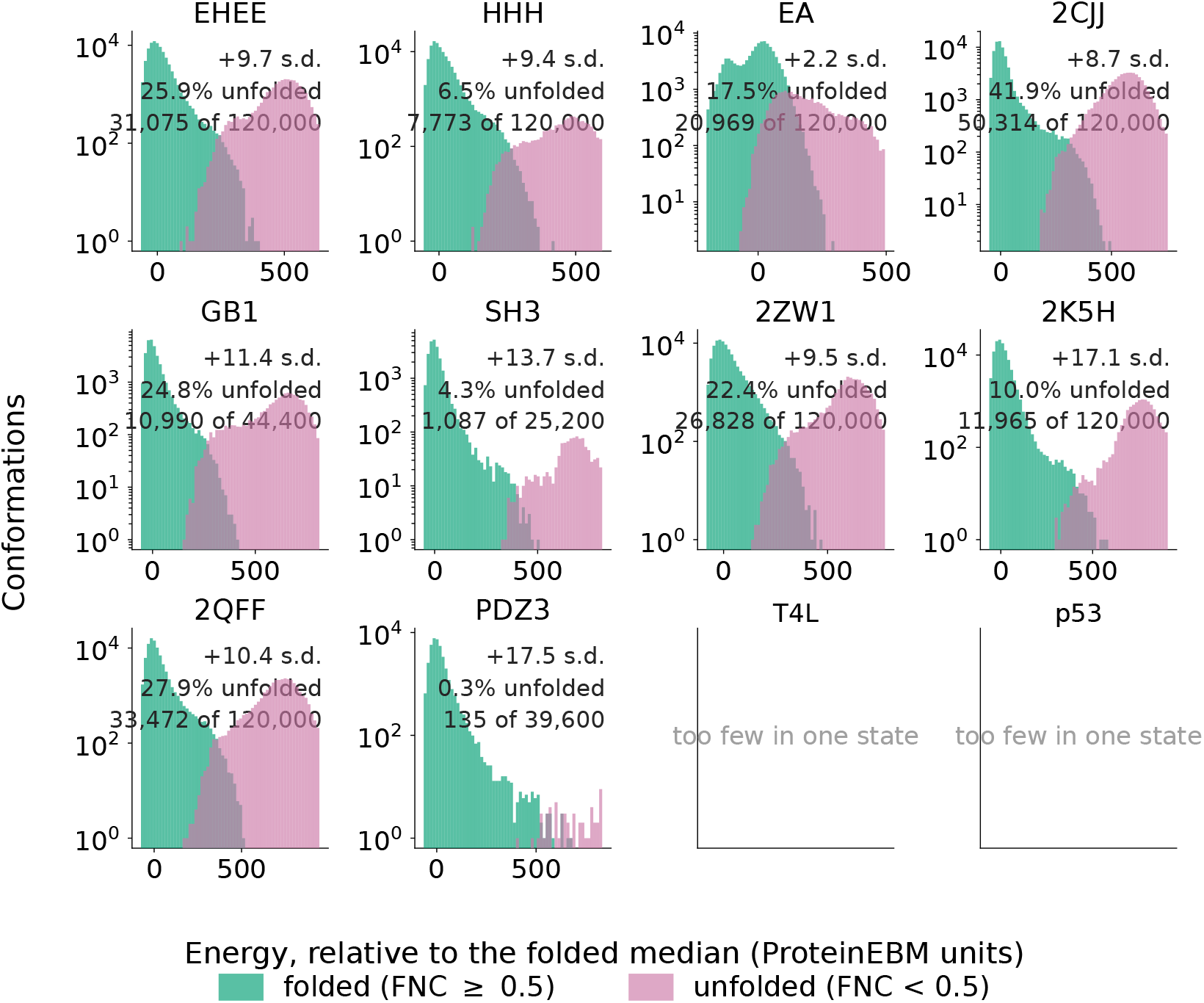
Per-panel distributions of ProteinEBM conformation energies, separated by BioEmu’s folded– unfolded classifier. For each of the twelve wild-type panels, histograms of per-conformation energies pooled across all variants of that panel. Blue, conformations classified as folded (fraction of native contacts ≥ 0.5); orange, conformations classified as unfolded (FNC *<* 0.5), using the same criterion BioEmu uses internally (Methods). Energies are reported in ProteinEBM units, shifted so that the folded-class median lies at zero on every panel. Annotations give the separation between class medians in units of the pooled within-class standard deviation, and the fraction of conformations in the unfolded class. Panels marked “too few in one state” (T4 lysozyme and p53 DBD) sample essentially only folded conformations, so the class separation is not defined. The figure shows that ProteinEBM assigns systematically higher energies to unfolded than to folded conformations of the same variant across all panels where both classes are populated, with class-median separations of 2 to 18 within-class standard deviations. The scoring readout in Fig. 3 takes the ensemble mean of these per-conformation energies as the per-variant ΔΔ*G*.

**Figure S4.**
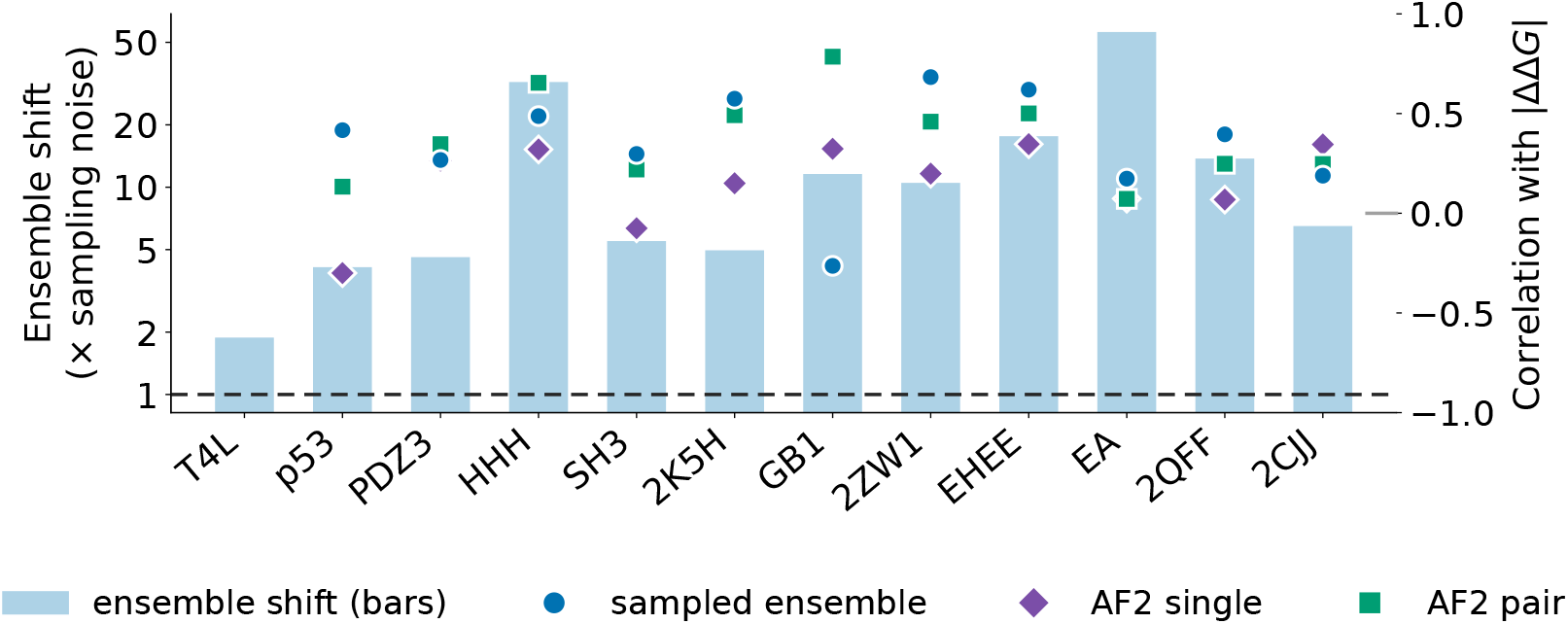
What a single substitution changes, and whether it tracks the effect. Bars, left axis: the median Jensen–Shannon divergence between a variant’s distribution of fraction of native contacts and the wild type’s, divided by the same divergence between two independent samplings of the wild type. A bar near one means the substitution moved the ensemble no more than resampling does. Markers, right axis: Spearman correlation between the size of each change and the magnitude of the measured effect. Diamonds, AlphaFold2 single representation; squares, its pair representation; circles, the sampled ensemble. The three are not a sequence — the two representations are produced together by the encoder and the ensemble is downstream of both — so the panel says which quantity carries more of the mutational effect. No correlation is shown for T4 lysozyme, whose seven mutants fall below the threshold of nine. Per-panel values and intervals are in Supplementary Table S1.

## Supplementary tables

**Table S1:**
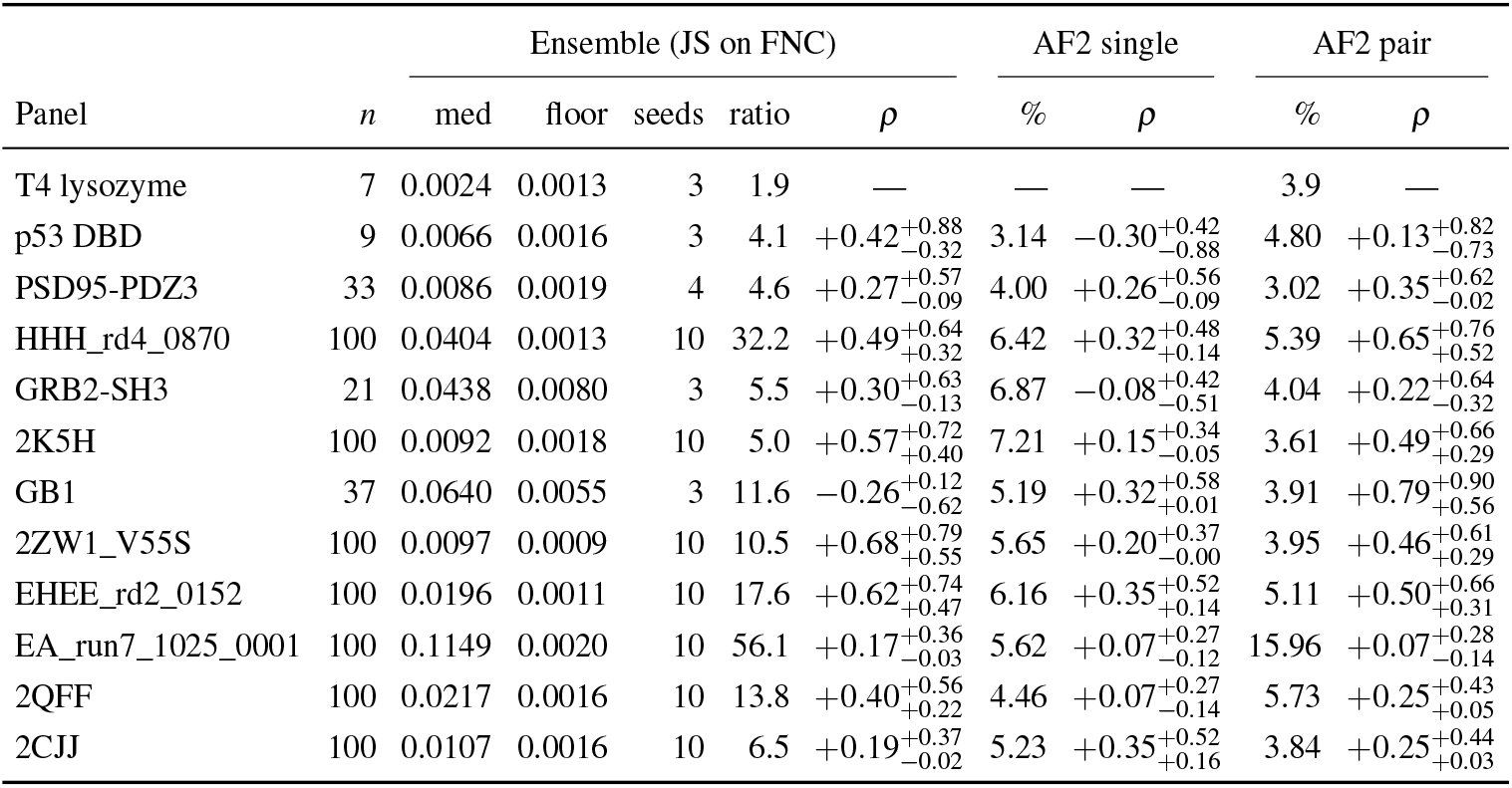
Ensemble and representation response to substitution, per panel. *n*, mutants. JS, median Jensen– Shannon divergence between a mutant’s distribution of fraction of native contacts and the wild type’s. Floor, the same divergence between two independent wild-type ensembles; seeds, the number of wild-type sampling seeds it rests on. Ratio, JS divided by the floor. Single % and pair %, median relative change in AlphaFold2’s single and pair representations under the substitution. Correlations are Spearman against the magnitude of the measured effect, with 95% percentile bootstrap intervals over variants, and are omitted below nine mutants. Both parts of the ratio are given because the floors differ almost fourfold across panels, so a ratio near one may arise from a small numerator or a large floor. Panels are ordered by the unfolded fraction BioEmu samples, as in Fig. 3b and Supplementary Fig. S4.

**Table S2:** Geometry of BioEmu’s folded and unfolded ensembles across benchmark panels. *N*_res_, chain length in residues. Folded and unfolded ensembles are classified per variant using BioEmu’s native-contact criterion. *R*_*g*_, median radius of gyration (Å). SD, standard deviation of *R*_*g*_ across sampled conformations (Å). Hofmann, median unfolded *R*_*g*_ divided by the empirical Flory scaling fitted to single-molecule scattering on 28 chemically denatured proteins [36]. AFRC, median unfolded *R*_*g*_ divided by the sequence-specific Analytical Flory Random Coil expectation [33–35]. Contact spec., long-range native-contact specificity relative to a compaction-matched self-avoiding chain. Panels where BioEmu samples virtually no unfolded conformations return missing values (—).

| Panel | $N_{\text{res}}$ | Folded | | Unfolded | | Ratio | | Contact spec. | |
| --- | --- | --- | --- | --- | --- | --- | --- | --- | --- |
| | | $R_g$ | SD | $R_g$ | SD | Hofmann | AFRC | folded | unfolded |
| EHEE_rd2_0152 | 40 | 9.30 | 0.16 | 12.84 | 3.19 | 0.73 | 0.81 | 4.73 | 1.45 |
| HHH_rd4_0870 | 43 | 9.38 | 0.26 | 13.37 | 3.31 | 0.73 | 0.81 | 7.53 | 1.76 |
| EA_run7_1025_0001 | 47 | 9.56 | 0.18 | 10.61 | 1.89 | 0.55 | 0.62 | 5.17 | 1.56 |
| 2CJJ | 54 | 10.56 | 0.33 | 14.74 | 3.53 | 0.70 | 0.80 | 10.75 | 1.84 |
| GRB2-SH3 | 56 | 9.80 | 0.14 | 13.38 | 2.98 | 0.63 | 0.72 | 4.79 | 2.06 |
| GB1 | 56 | 10.29 | 0.10 | 14.00 | 3.57 | 0.65 | 0.75 | 5.51 | 1.55 |
| 2ZW1_V55S | 57 | 10.41 | 0.14 | 13.48 | 2.81 | 0.62 | 0.72 | 5.29 | 1.56 |
| 2K5H | 62 | 10.09 | 0.25 | 14.52 | 2.97 | 0.64 | 0.74 | 4.93 | 1.23 |
| 2QFF | 70 | 11.84 | 0.18 | 15.33 | 3.47 | 0.63 | 0.73 | 10.35 | 1.80 |
| PSD95-PDZ3 | 84 | 11.44 | 0.11 | — | — | — | — | — | — |
| T4 lysozyme | 164 | 16.40 | 0.46 | — | — | — | — | — | — |
| p53 DBD | 194 | 16.39 | 0.21 | — | — | — | — | — | — |

